# Mass evacuation and increases in long-term care benefits: lessons from the Fukushima nuclear disaster

**DOI:** 10.1101/669499

**Authors:** Tomohiro Morita, Michihito Ando, Yui Ohtsu

**Author notes:** **Corresponding author:** Tomohiro Morita, Soma Central Hospital, 3-5-18, Okinouchi, Soma City, Fukushima, 976-0016.

## Abstract

**Background:** Though mass evacuation may increase the need for long-term care (LTC) services, how the need for LTC services increases and how the public LTC system affects it is not well understood. We evaluated changes in public LTC benefits for the people living in the mandatory evacuation areas established after the 2011 Fukushima nuclear disaster and examined the roles of the universal LTC insurance system in Japan.

**Methods:** In order to evaluate the effect of the mandatory evacuation on LTC benefits, we examined the trends of LTC benefits in the Fukushima evacuation group and the nationwide non-evacuation group. We first decomposed per-elderly-individual benefits at the municipality level into the LTC certification rate and per-certified-individual benefits, and then implemented difference-in-differences analysis using these variables as outcomes.

**Results:** Per-elderly-individual benefits significantly increased from 2012 onward in the evacuation group, and this was explained by an increase in the certification rate rather than in per-certified-individual benefits. Increases in per-elderly-individual benefits and the certification rate in the post-disaster period were observed in all but the highest care level, and the corresponding outcomes for the highest care level decreased immediately after the disaster. We also found that the increase in the certification rate had been mostly realized by an increase in the number of certified individuals.

**Conclusions:** The increase in LTC benefits can be associated with the impact of the increase in the number of people newly certified to receive LTC benefits after the mandatory evacuation. In order to cope with the increase in utilization of long-term care and associated costs after disasters in aging societies, both formal long-term care services and social support for informal care for evacuees should be considered important.

## 1. Introduction

Natural and manmade disasters, such as earthquakes, floods, and fires, often lead to mass evacuation and/or relocation. In 2015, there were 27.8 million people newly involved in evacuation or relocation caused by disasters worldwide [1]. Damage to communities following a mass evacuation has become a social issue. As the number of elderly people increases throughout the world, the health effects due to damage to communities seem likely to become more severe, but these adverse effects, especially in the long-term, are not clearly understood.

Mass evacuation in an aging society is exemplified by the case of the 2011 Fukushima nuclear disaster, which occurred after the Great East Japan Earthquake and Tsunami on March 11, 2011. Following a series of government evacuation orders, more than 160,000 citizens were evacuated from the areas surrounding the nuclear power plant [2]. In particular, the entire population of the nine municipalities closest to the damaged nuclear plant, nearly 80,000 citizens, was forced to evacuate.

This mass evacuation not only raised the mortality of the elderly in the short term [3], but may also have increased the utilization of public long-term care (LTC) services [4–6]. However, *how* such an evacuation affects the utilization of LTC services is still not well understood. Do the elderly people who had already been using LTC services before the evacuation use more LTC services after it? Or do the elderly people who had not been using any LTC services before the evacuation start doing so after they have been evacuated? The objective of this study is to assess the factors behind the increase in LTC utilization after an evacuation, which are important for policy makers in countries that frequently experience natural disasters and subsequent evacuations. We examined how the utilization of public LTC benefits changed after the evacuation among citizens who had lived in the mandatory evacuation areas of the 2011 Fukushima nuclear disaster using the difference-in-differences (DID) method.

## 2. Materials and methods

### 2.1. Settings

After the 2011 Fukushima nuclear disaster, evacuation instructions were issued by the Japanese government on March 12, 2011 for a 20km area surrounding the nuclear plant and in areas where the annual cumulative radiation dose was expected to exceed 20 mSv/year. As a result, eight whole municipalities (the villages of Iitate, Katsurao, and Kawauchi and the towns of Namie, Futaba, Okuma, Tomioka, and Naraha) were designated as mandatory evacuation areas. One additional municipality, the town of Hirono, also decided to issue a mandatory evacuation order to all of its residents at the direction of its mayor. As a result, a total of 78,768 residents were forced to evacuate. According to the National Population Census, elderly residents (aged 65 and older) accounted for 25.1% (19,792/78,768) of the population in these nine municipalities at the time of the disaster. Parts of three municipalities (the cities of Minamisoma and Tamura and the town of Kawamata) were also designated as mandatory evacuation areas. In total, 164,865 Fukushima residents were evacuated.

In Japan, all people aged 65 and older have been insured by public LTC insurance since 2000. When an individual wants to make use of this insurance, municipalities conduct interviews and surveys concerning his or her living situation. An initial assessment is then used to assign the applicant to one of the eight LTC need levels (not certified, support level 1–2, and care level 1–5). The final LTC need level is decided by the Care Needs Certification Board, a committee of medical and other professionals. The maximum amount of expenditure covered by the LTC insurance is fixed in accordance with the applicant’s level as determined by this process [7].

### 2.2. Data

In order to evaluate the effect of the mandatory evacuation on LTC benefits, we examined the LTC benefits in the eight “evacuation” municipalities and compared them to those observed nationwide.[8]

We first define *Q*_*it*_ as the aggregated quantity of LTC benefits, which is measured as the amount of LTC benefits per elderly individual for municipality *i* in fiscal year *t*. Then we can decompose *Q*_*it*_ as follows [9]:

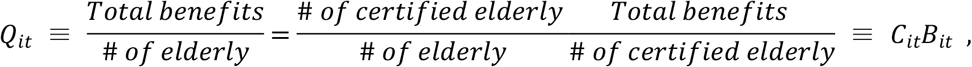

where *C*_*it*_ is the LTC certification rate (i.e. the ratio of elderly people certified to receive LTC benefits to the total elderly population in question) and *B*_*it*_ is the amount of LTC benefits per certified individual. We call *Q* “per-elderly-individual benefits”, *C* “certification rate”, and *B* “per-certified-individual benefits” and examine these three variables.

In addition, because the certified elderly are categorized into seven care levels (support level 1-2 and care level 1–5), we also investigate LTC benefits disaggregated by care level. For example, we use disaggregated outcomes for care level 1 as

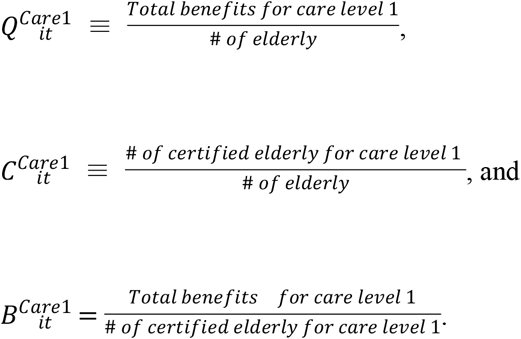

We can also define disaggregated outcomes for support levels 1-2 and care levels 2-5 in the same manner. See Table 1 for the descriptive statistics of the aggregated and disaggregated outcome variables. All the variables are constructed from the Status Report on the Long-term Care Insurance collected by the Ministry of Health, Labour and Welfare of Japan. This report provides municipality-level data based on administrative data about all the benefits dispensed through public LTC insurance for each municipality in an open public database. From this public database, we extracted the annual data on LTC benefits, the number of elderly people certified to receive LTC benefits, and the population over 65 years of age in each municipality. Some municipalities organized or joined inter-municipality insurance coalitions, and some municipalities merged during the sample period (2007-2014). In these cases, we aggregated the LTC benefits data and population data based on municipalities as they were constituted in 2014.

**Table 1:**
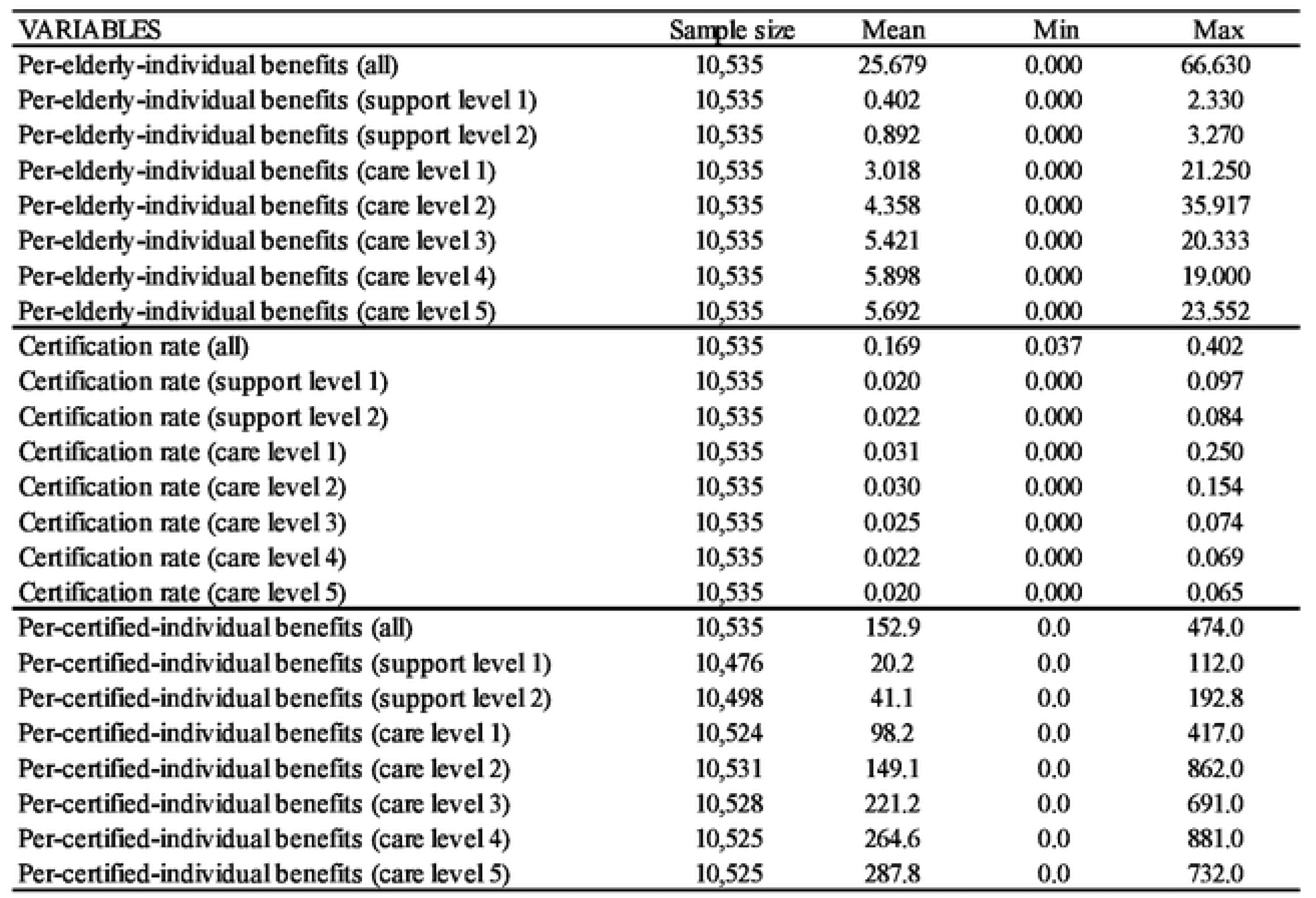
Descriptive statistics. Note that data for per-certified-individual benefits in some care levels arc missing when the number of certified individuals in these care levels arc is zero. In addition, the maximum value of per-certified-individual benefits for care level 2 appears to be unusually high, but this is due to the fact that this variable is calculated as the ratio of annual LTC benefits to the number of certified elderly individuals at a certain point. Thus this variable tends to fluctuate in municipalities with small numbers of certified elderly individuals. Source: The Survey of Long-term Care Benefit Expenditures. The unit of “benefit” is 1,000 points (i.e. 10,000 JPY or around 100 USD in standard areas).

When it comes to the number of certified elderly individuals and the population over 65 years of age, these values are measured in the last month of the fiscal year (March of the following calendar year because the Japanese fiscal year starts in April). LTC benefits, which are measured in so-called “benefit points” (1 point = 10 yen in a standard area), are in turn aggregated from March to February of the following calendar year. This is somewhat confusing because the first month is March, not April (the first month of the fiscal year). This is advantageous in our case, however, because the disaster occurred on the 11^th^ of March, 2011, and therefore the survey of the year for 2011 consists of data concerning only post-disaster months (March 2011-Februrary 2012).

It should be noted that the residential status (registered place of residence) of citizens remained the same after the mandatory evacuation if they wanted it to, and they were given financial support unless they changed it [10]. Most of the LTC benefits for these citizens were therefore financed by these nine municipalities before, during, and after the evacuation. These circumstances enable us to assess the impact of the mandatory evacuation on LTC benefits.

### 2.3. Methods

In order to investigate how this evacuation has affected the utilization of LTC services, we examine how per-elderly-individual benefits (*Q*), certification rate (*C*), and per-certified-individual benefits (*B*) have changed over time in the evacuation group (i.e. those who had been living in evacuation areas before the disaster) by implementing simple difference-in-differences (DID) estimations. The estimation model is expressed as follows:

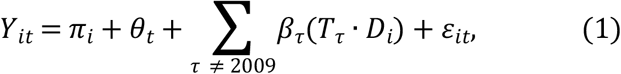

where *Y*_*it*_ is either *Q*_*it*_, *C*_*it*_, or *B*_*it*_, *π*_*i*_ is a municipality fixed effect, *θ*_*t*_ is a time fixed effect, *T*_*τ*_ is a time dummy variable that takes one if *t* = *τ*, *D*_*i*_ is a “treatment” dummy variable that takes one if municipality *i* is entirely included in evacuation areas in Fukushima, and *ε*_*it*_ is a random error.

As explained in Section 2.1, the entirety of each of the eight municipalities is included in the “evacuation areas” and one municipality issued a mandatory evacuation order to all of its residents, resulting in nine “treated” municipalities in our sample. Note that we exclude municipalities with partial evacuation orders or without a mandatory evacuation order in the Fukushima region (Fukushima prefecture) and in tsunami-affected coastal areas from the “control” group (*D*_*i*_ = 0): these municipalities presumably have been affected by the nuclear disaster and/or tsunami in 2011 and are not suitable for constructing the counterfactual trends to the evacuation areas.

The parameter of interest is *β*_*τ*_, which reflects a differential trend in the outcome variable for the treated (evacuation) group at *t* = *τ*, compared with a corresponding trend for the control (non-evacuation) group. Because we exclude the time dummy variable at *t* = 2009, *β*_*τ*_ captures E(*Y*_*iτ*_ − *Y*_*i*2009_|*D*_*i*_ = 1) − E(*Y*_*iτ*_ − *Y*_*i*2009_|*D*_*i*_ = 0), which is identical to a conventional DID parameter. Note that fiscal year 2009 is the latest pre-disaster year when the complete annual data is available for the evacuation group. That is, the data for fiscal year 2010 is not available because the data for February 2011, which was supposed to be included in fiscal year 2010, is not available due to administrative disorder during the evacuation period (the Fukushima nuclear disaster occurred in March 2011).

As is now common in DID and event study literature [11, 12], in the pre-disaster period, *τ* < 2009, *β*_*τ*_ serves as a placebo parameter whose estimated value should be near zero if no differential trends exist between treated and control municipalities. In the post-disaster period, *τ* ≥ 2011, *β*_*τ*_ captures how the trend of mean outcomes for the treated municipalities deviates from the trend for the control municipalities given that no differential trends exist in the pre-disaster period.

Before examining estimation results, Figure 1 shows the trends of outcome variables on average for the treated group and the control group. These graphs indicate that in the pre-disaster period (2007-2009) the trends were more or less similar between the evacuation and control groups, indirectly validating the common trend assumption of DID estimation. In the post-disaster period (2011-2014) they significantly differed. In 2011, the year of the disaster and evacuation, the certification rate (*C*) sharply increased in the evacuation group, whereas per-certified-individual benefits (*B*) sharply decreased, resulting in a modest increase in per-elderly-individual benefits (*Q*). Since 2012, the certification rate (*C*) has remained high, and per-certified-individual benefits (*B*) have more or less recovered to their pre-disaster trend. As a result, per-elderly-individual benefits (*Q*) have sharply increased.

**Figure 1:**
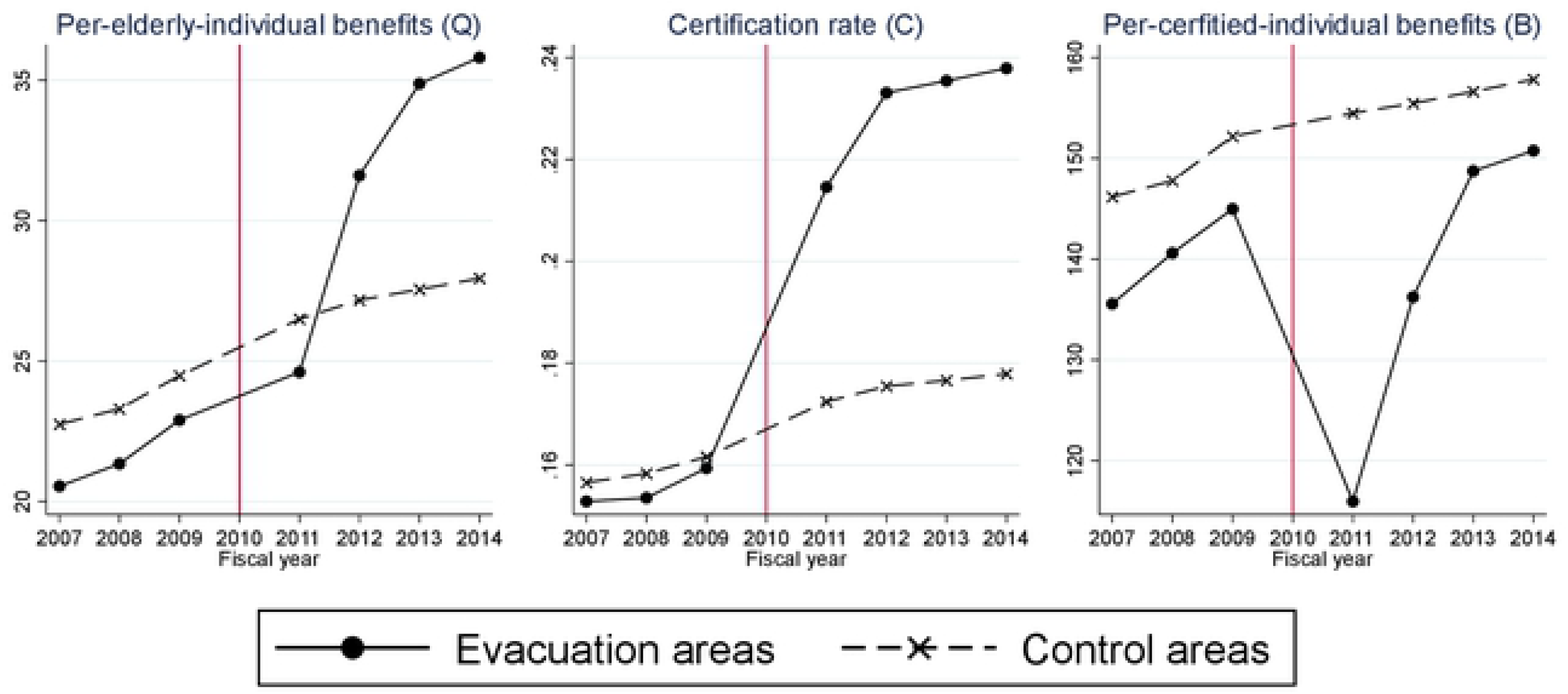
Time trends of long-term care benefits in the Fukushima mandatory evacuation group and the non-evacuation control group. Per-elderly-individual benefits: long-term-care benefits per elderly person. Certification rate: the percentage of people aged 65 and older who were certified to receive long-term care services. Per-certified-individual benefits: long-term care benefits per certified individual. The unit of “benefit” is 1,000 points (10,000 JPY or around 100 USD in standard areas).

## 3. Results

Figure 2 presents DID estimation results based on equation (1). The three graphs in this figure show estimation results using different aggregated outcomes, namely per-elderly-individual benefits (*Q*), certification rate (*C*), and per-certified-individual benefits (*B*) respectively. These outcome variables are based on all LTC benefits and ertified individuals.

**Figure 2:**
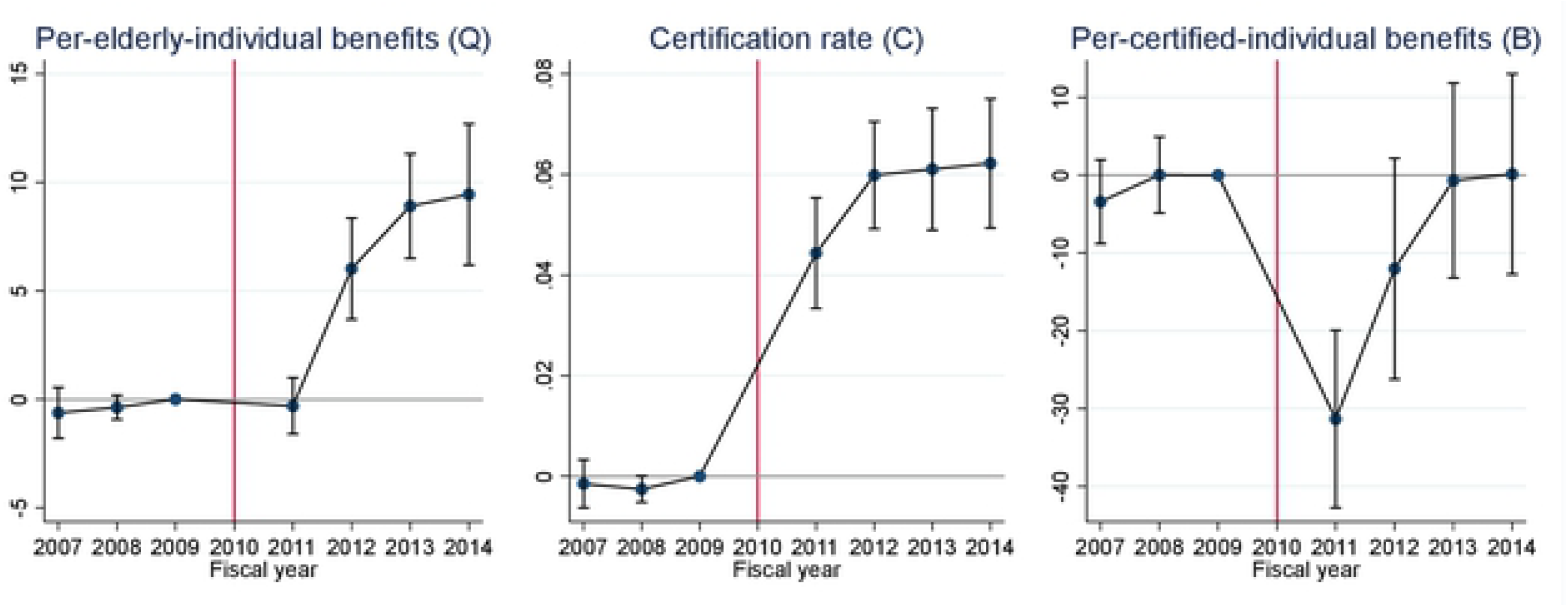
Difference-in-differences (DID) estimates and their 95% confidence intervals. Confidence intervals are calculated by clustered robust standard errors clustered by municipality. All the DID estimates are estimates for *β*_*τ*_ in equation (1). The unit of “benefit” is 1,000 points (10,000 JPY or around 100 USD in standard areas).

The results of this analysis imply that per-elderly-individual benefits (*Q*, left graph) increased from 2012 onward, and that this was explained by an increase in the certification rate (*C*, center graph) rather than per-certified-individual benefits (*B*, right graph). DID estimates for the certification rate are positive and statistically significant immediately after the evacuation, and the magnitude of the estimates implies that the evacuation has increased the certification rate by around 6 percentage points. DID estimates for per-certified-individual benefits decreased sharply in 2011 (and modestly in 2012), and this in turn is presumably a result of the disaster (earthquake and tsunami) and the ensuing evacuation necessitated by the nuclear plant accident. Note also that placebo estimates in the pre-disaster period are around zero for all three outcomes, suggesting that the “parallel trends” assumption of DID seems to be plausible in the post-disaster period.

Figure 3 shows DID estimates for disaggregated outcomes by care level (see appendix Figure A.1-A.2 for the same estimation results with separated graphs). Firstly, the results of this analysis present a markedly different pattern in DID estimates for care level 5 outcomes. Negative estimates of the care level 5 certification rate (*C*) after 2011 suggest that some elderly people in this category may not have survived the disasters and evacuation (center graph). The estimates in this category gradually approach zero from 2011 to 2014, however, implying that the care level 5 certification rate was returning to its pre-evacuation level. DID estimates for care level 5 per-certified-individual benefits (*B*) are also strongly negative in 2011 and 2012. This is probably due to the evacuation, although the placebo estimates in 2008 fluctuate unstably (right graph). The estimates in this category also approached zero in 2013 and 2014.

**Figure 3:**
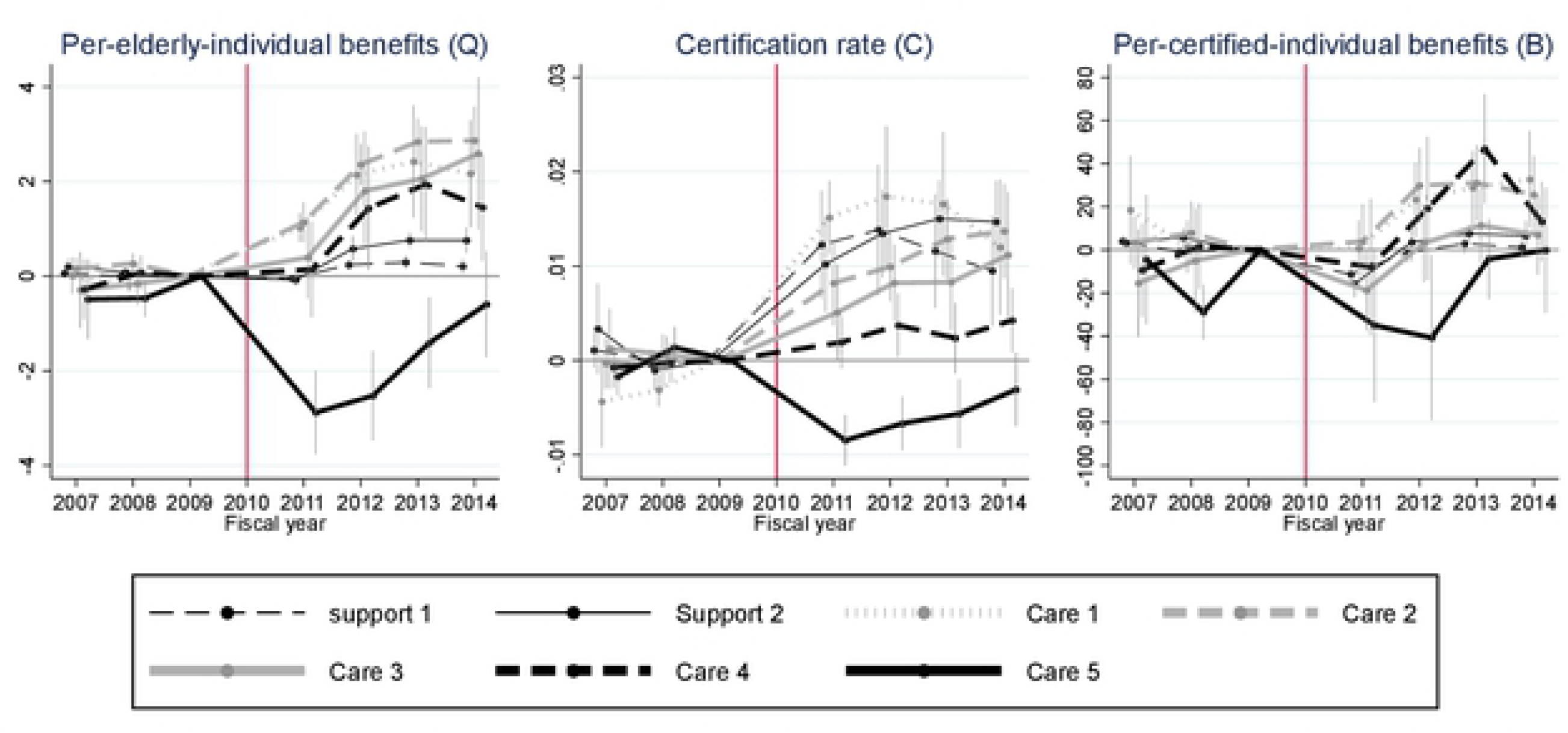
Difference-in-differences (DID) estimates for each care level and their 95% confidence intervals. Confidence intervals are calculated by clustered robust standard errors clustered by municipality. All DID estimates are estimates for *β*_*τ*_ in equation (1). The unit of “benefit” is 1,000 points (10,000 JPY or around 100 USD in standard areas).

Secondly, when it comes to DID estimates for support levels 1-2 and care levels 1-4 in Figure 3, these show more or less similar trends that are consistent with the DID results in Figure 2. There are, however, two points worth mentioning. First, for these support/care levels, DID estimates for per-elderly-individual benefits (*Q*) and the certification rate (*C*) are always positive and almost always significantly different from zero from 2012 onward (left and center graphs. See 95% C.I. in appendix Figure A.1. and A.2.). The contribution to the aggregated positive impact on per-elderly individual benefits is higher in care levels 1-3 than the other levels, but the DID estimates for the certification rate suggest that the number of certified individuals has been sharply rising in all care-need categories but care levels 4 and 5. Another interesting finding is that DID estimates for per-certified-individual benefits (*B*) are also positive for 2012-2014 (right graph), although the 95% C.I. is consistently above zero only for care levels 1 and 2 (appendix Figure A.1, A.2).

In summary, the Fukushima disaster and the ensuing evacuation increased the overall certification rate but not overall per-certified-individual benefits (Figure 2). If we look at disaggregated outcomes by care level, the disaster and evacuation decreased the certification rate and per-certified-individual benefits for the highest care level, and increased the certification rates for the other care levels and the per-certified-individual benefits for some care levels (Figure 3).

Finally, given the finding that the positive impact of the evacuation on LTC benefits was mainly owing to an increase in the certification rate, it is worth looking into what drove this increase in this group. We therefore compared the trends of the numerator and the denominator of the certification rate (i.e. the number of certified people and the total number of elderly people) between the evacuation and the control groups. Figure 4 clearly shows the main source of the increase in the certification rate in the evacuation group is an increase in the number of certified elderly (the numerator), although the number of those aged 65+ (the denominator) also decreased modestly in 2011.

**Figure 4:**
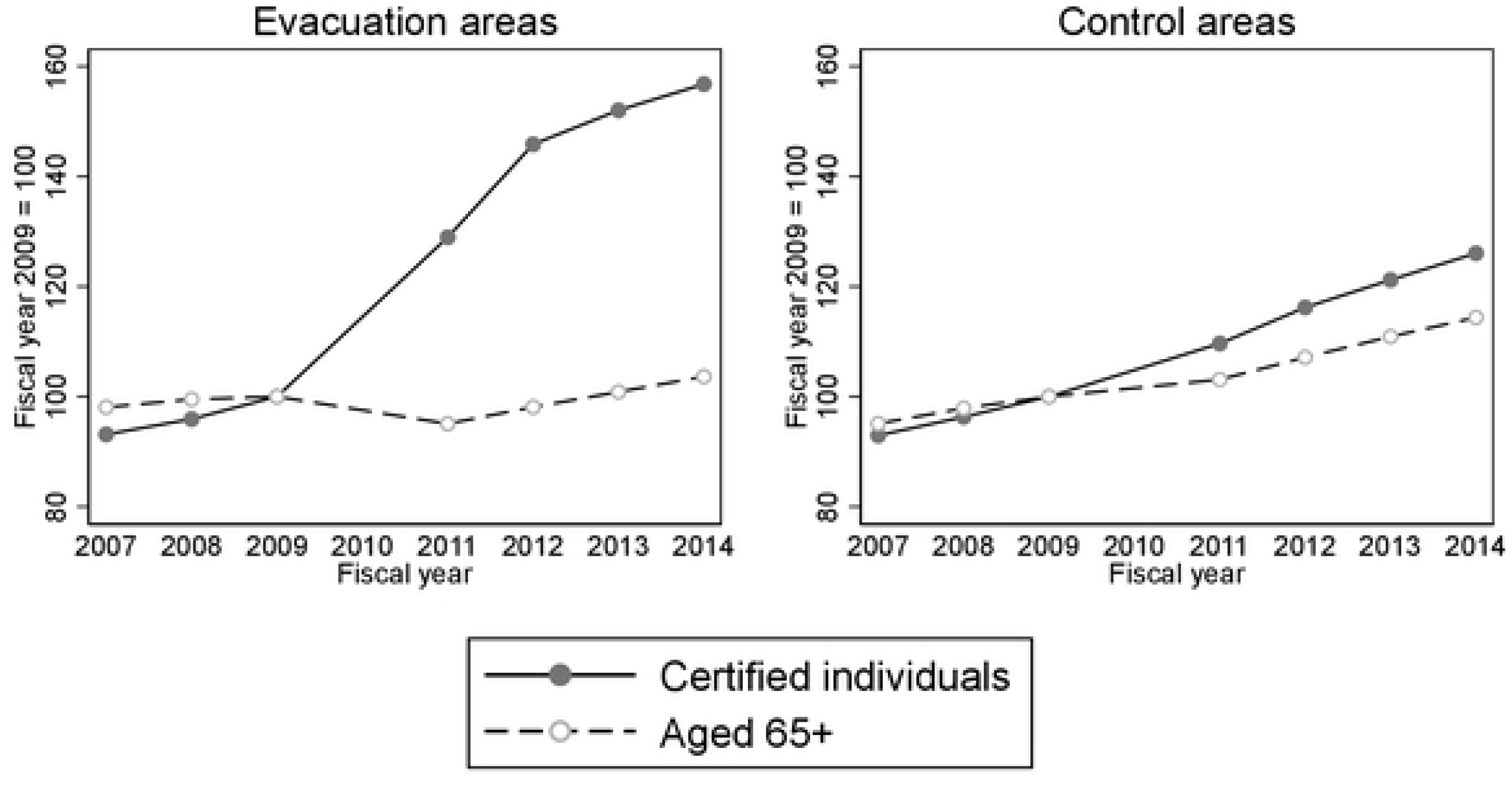
Trends in the numbers of certified individuals and aged 65+ in the evacuation and control areas. Certified individuals: the number of people aged 65 and older who were certified to receive long-term care services. Aged 65+: the number of people aged 65 and older. The numbers are standardized to 100 in 2009.

## 4. Robustness checks

Figure 2 clearly shows that the increase in LTC benefits per elderly individual after the disaster is caused by the increase in the certification rate, not in LTC benefits per certified individual. In this section we provide the results of several robustness checks on these primary findings. We implement robustness checks by estimating the same equation (1) using different combinations of the following two estimation settings.

First, we use two different weighting schemes for estimation: (1) not weighting observations as in our baseline estimation (i.e. ordinary least squares regression: OLS) and (2) weighting observations by the number of elderly individuals for the outcomes of *Q* and *C* and by the number of certified individuals for the outcome of *B* (i.e. weighted least squares regression: WLS). There are pros and cons for weighting observations in aggregated data, but our purpose in using different weights is to ensure that our main findings are robust to different weighting schemes [13].

Second, we check the sensitivity of DID estimates by using three different samples (i.e. three different control groups). In the baseline estimation we used all the Japanese municipalities except for partial-evacuation and non-evacuation municipalities in Fukushima prefecture and tsunami-affected coastal areas. In this section we use the following three different samples: the same municipalities as in the baseline estimation (the baseline sample), the municipalities in the baseline sample whose outcome values (*all* of *Q*, *C* and *B*) are within the minimum and maximum of the outcome values of the treated municipalities in all the three pre-disaster years (the trimmed sample), and evacuation municipalities and non-evacuation municipalities in Fukushima prefecture (the Fukushima sample).

In the case of the trimmed sample, 399 out of 1,496 control municipalities in the baseline sample are kept in the sample. This trimming procedure is somewhat arbitrary, but it enables us to exclude municipalities whose outcome values are far away from and not comparable to those of the evacuation municipalities in the pre-disaster period. Appendix Figure A.3 shows that this trimming procedure does in fact make the averaged outcomes of the control areas quite similar to those of the evacuation areas in the pre-disaster period.

In the Fukushima sample, we use 47 non-evacuation municipalities in Fukushima, which are excluded from the baseline sample, as an alternative control group. We keep partial-evacuation municipalities in Fukushima out of the control group. As we already discussed, non-evacuation municipalities in Fukushima may not be an appropriate control group. It is nevertheless worthwhile to compare the evacuation municipalities in Fukushima with the non-evacuation municipalities in Fukushima because of their geographic and socio-economic similarities in the pre-disaster period.

Figure 5 provides the results of our robustness checks. Because we use two weighting schemes (OLS and WLS) and three samples (baseline, trimmed, and Fukushima samples), we implemented six, including one baseline, DID estimations for each outcome. Some estimated coefficients are different from the baseline DID estimation (i.e. OLS with the baseline sample) but overall tendencies are similar to the baseline analysis for all three outcomes.

**Figure 5:**
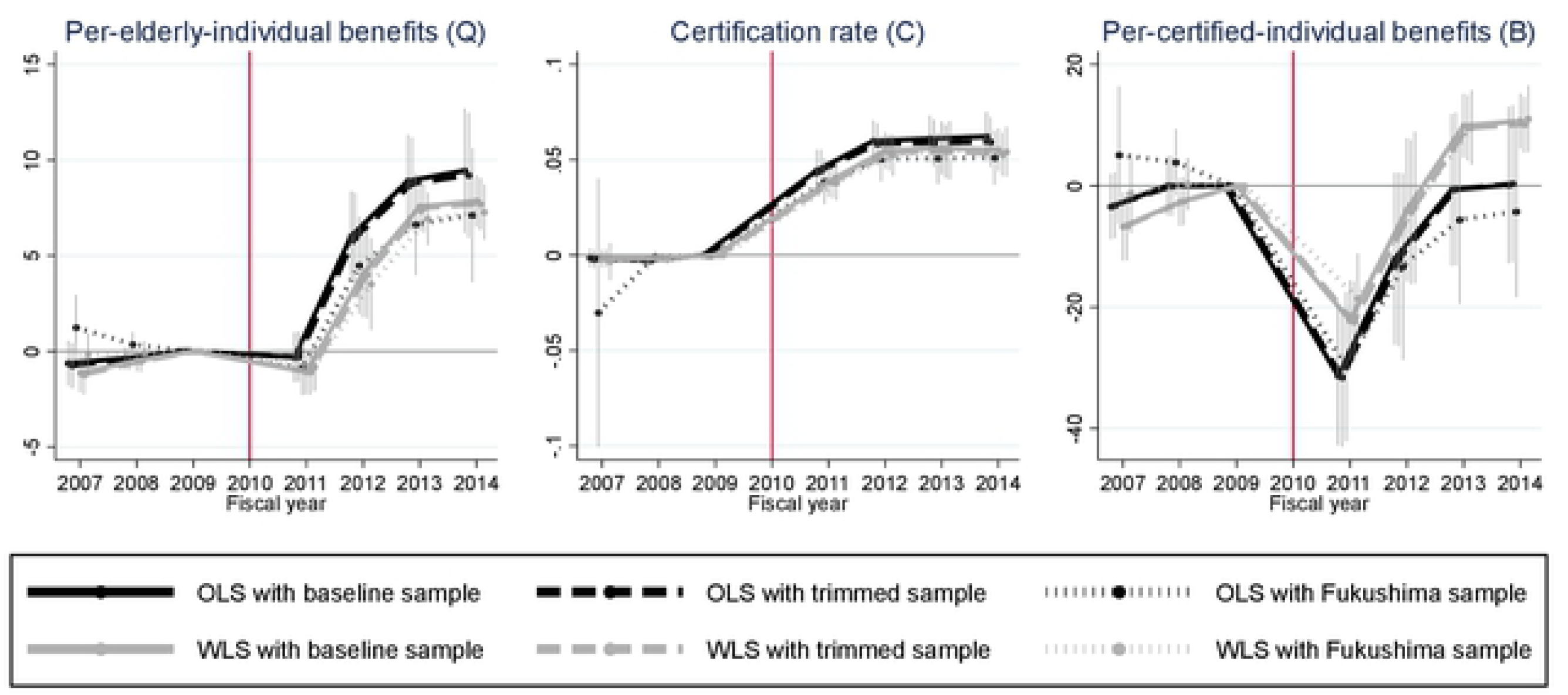
Difference-in-differences (DID) estimates for different weighting schemes and samples and their 95% confidence intervals. Confidence intervals are calculated by clustered robust standard errors clustered by municipality. All DID estimates are estimates for *β*_*τ*_ in equation (1). The unit of “benefit” is 1,000 points (10,000 JPY or around 100 USD in standard areas). “OLS” is a normal ordinary least squares regression and “WLS” is a weighted least squares regresssion in which the weight is the number of elderly individuals for the outcomes of *Q* and *C* and the number of certified individuals for the outcome of *B*. The trimmed sample includes the municipalities in the baseline sample whose outcome values are within the minimum and maximum of outcome values of the treated municipalities in all three pre-disaster years. The Fukushima sample contains the evacuation and non-evacuation municipalities in Fukushima prefecture.

## 5. Discussion

It has been reported that LTC benefits have increased in the peripheral area of the damaged nuclear plant [14]. Our findings suggest that the increase in LTC benefits can be associated with the impact of the increase in the number of people newly certified to receive LTC benefits after the mandatory evacuation.

There are several reasons for this. First, as the communities in the mandatory evacuation zones were severely damaged, the amount of informal care provided by young members of the community seems to have declined after the disaster. Second, evacuation could have worsened ADL, dementia, and other health problems among the elderly population [15]. Our study was consistent with previous studies demonstrating that the number of people relying on public LTC increased after the disaster [4–6]. Third, residents in the mandatory evacuation areas were exempted from copayment according to the compensation program. This financial support could have increased the demand for long-term care [16].

In addition, it is possible that with their greater mobility and resilience the healthy elderly population in a community who do not require long-term care may be more likely than the rest of the population to change their resident registration from the evacuation areas to other areas even though it may lead to the cessation of financial support. Our analysis, however, found only a modest decline in the number of elderly people in the evacuation group.

Another important finding is that both the certification rate and the per-certified-individual benefits for the highest care level (care level 5) decreased immediately after the disaster (2011), while the corresponding outcomes for the lower care levels tended to gradually increase after the evacuation. Such heterogeneous effects on the evacuated elderly suggest that special care soon after the disaster is particularly important for the most vulnerable group, whereas increasing LTC needs among the elderly due to the evacuation are of greater importance from a longer perspective.

This study demonstrates the possibility that an increase in the utilization of LTC services immediately after a mass evacuation can be attributed to increases in people newly certified to receive LTC benefits. This implies that universally provided LTC insurance might have worked as a quick buffer that mitigates some of the negative impacts of evacuation, such as the loss of informal care and evacuation-related health deterioration [17]. At the same time, it highlights the importance of the question of how such exceptional costs incurred by evacuations are to be financed as a policy issue to be addressed in the context of the current municipality-based financing system of Japanese LTC insurance. In order to cope with the increase in need for long-term care and associated costs after disasters in aging societies, both formal long-term care services and social support for informal care for evacuees should be considered important.

## APPENDICES

**Figure A.1:**
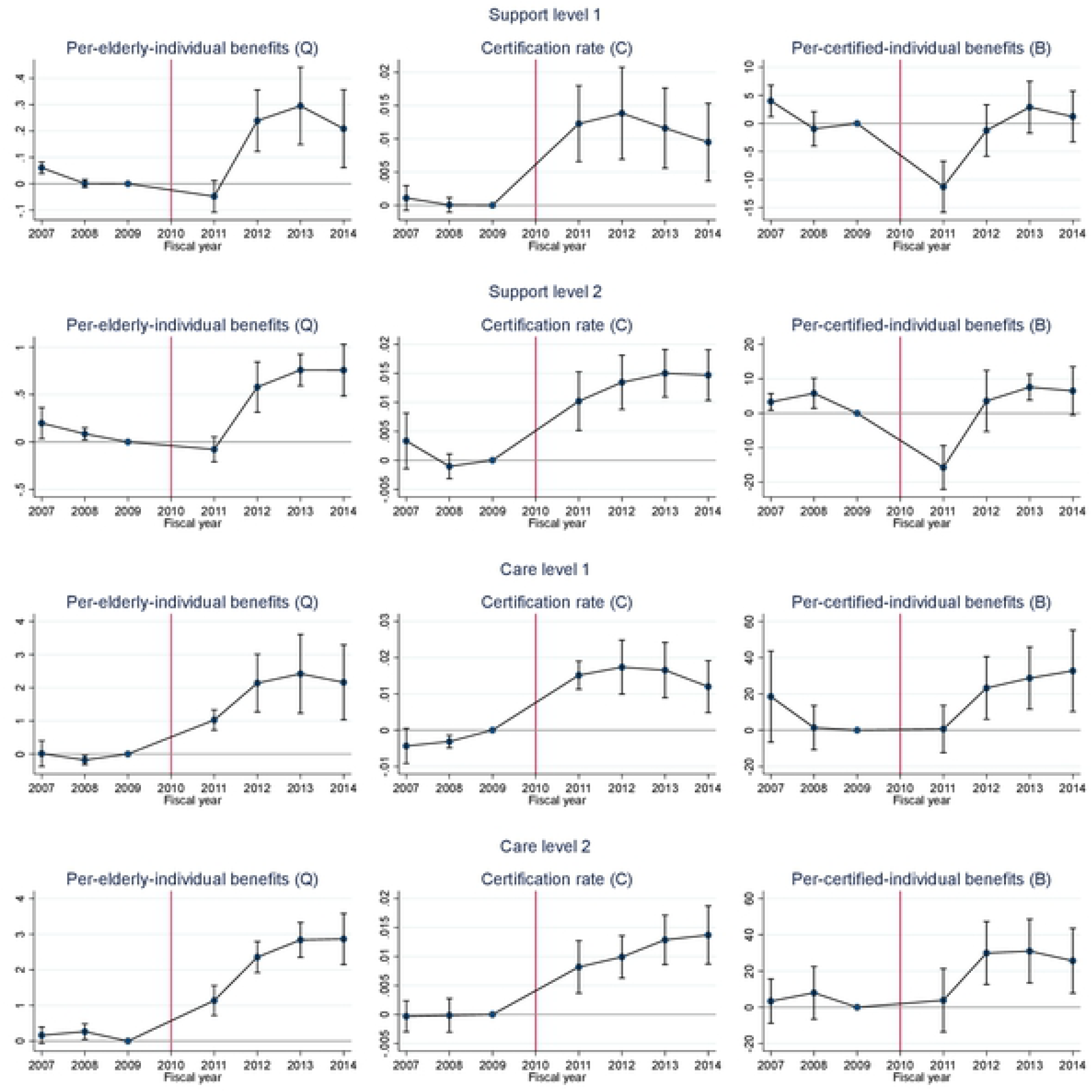
Difference-in-differences (DID) estimates and their 95% confidence intervals (lower care levels). Confidence intervals are calculated by clustered robust standard errors clustered by municipality. All DID estimates are estimates for *β*_*τ*_ in equation (1). The unit of “benefit” is 1,000 points (10,000 JPY or around 100 USD in standard areas).

**Figure A.2:**
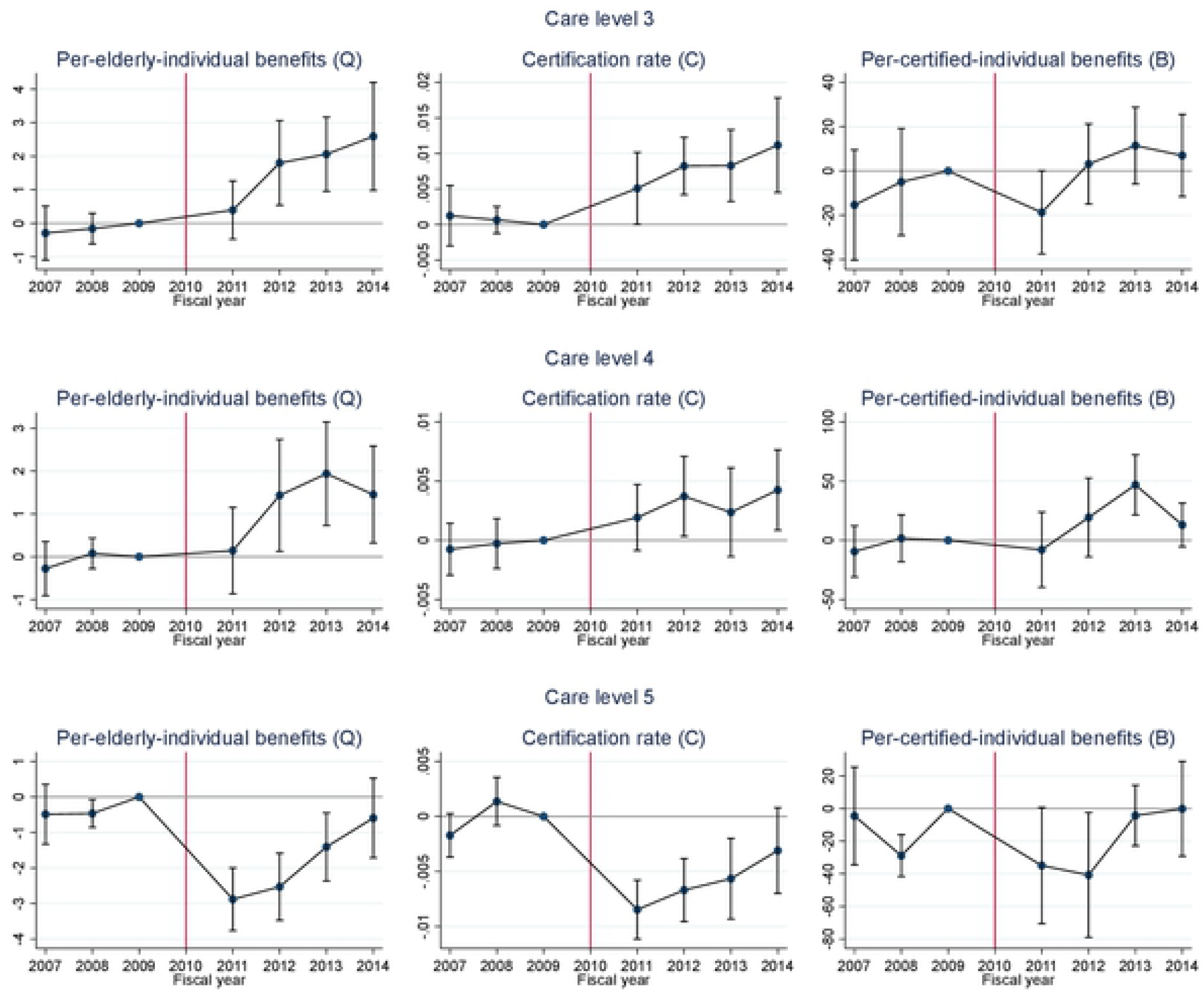
Difference-in-differences (DID) estimates and their 95% confidence intervals (higher care levels). Confidence intervals are calculated by clustered robust standard errors clustered by municipality. All DID estimates are estimates for *β*_*τ*_ in equation (1). The unit of “benefit” is 1,000 points (10,000 JPY or around 100 USD in standard areas).

**Figure A.3:**
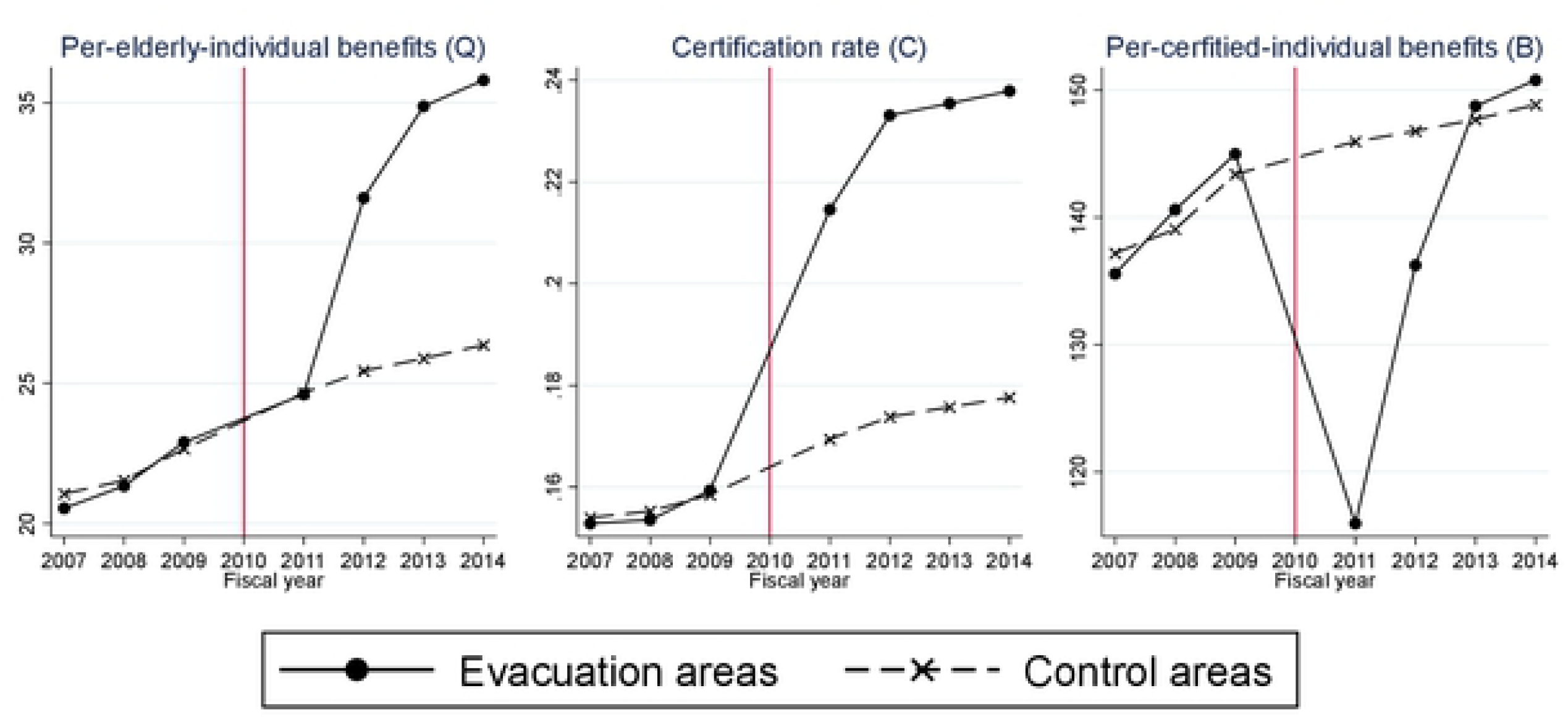
Time trends of long-term care benefits in the Fukushima mandatory evacuation group and the non-evacuation control group with the trimmed sample. Per-elderly-individual benefits: long-term-care benefits per elderly person. Certification rate: the percentage of people aged 65 and Ider who were certified to receive long-term care services. Per-certified-individual benefits: long-term care benefits per certified individual, The unit of “benefit” is 1,000 points (10,000 JPY or around 100 USD in standard areas). The trimmed sample inncludes the municipalities in the baseline sample whose outcome values are within the minimum and maximum of outcome values of the treated municipalities in all three pre-disaster years.

